# The thin edge of the wedge: extremely high extinction risk in wedgefishes and giant guitarfishes

**DOI:** 10.1101/595462

**Authors:** Peter M. Kyne, Rima W. Jabado, Cassandra L. Rigby, Dharmadi, Mauvis A. Gore, Caroline M. Pollock, Katelyn B. Herman, Jessica Cheok, David A. Ebert, Colin A. Simpfendorfer, Nicholas K. Dulvy

## Abstract

1. The process of understanding the rapid global decline of sawfishes (Pristidae) has revealed great concern for their relatives, the wedgefishes (Rhinidae) and giant guitarfishes (Glaucostegidae), not least because all three families are targeted for their high-value and internationally-traded ‘white’ fins.
2. The objective of this study was to assess the extinction risk of all 10 wedgefishes and 6 giant guitarfishes by applying the International Union for Conservation of Nature (IUCN) Red List Categories and Criteria, and to summarise their biogeography and habitat, life history, exploitation, use and trade, and population status.
3. Wedgefishes and giant guitarfishes have overtaken sawfishes as the most imperilled marine fish families globally, with all but one of the 16 species facing an extremely high risk of extinction due to a combination of traits – limited biological productivity, presence in shallow waters overlapping with some of the most intense and increasing coastal fisheries in the world, and over-exploitation in target and bycatch fisheries driven by the need for animal protein and food security in coastal communities and trade in meat and high-value fins.
4. Two species with very restricted ranges, the Clown Wedgefish (*Rhynchobatus cooki*) of the Indo-Malay Archipelago and the False Shark Ray (*Rhynchorhina mauritaniensis*) of Mauritania may be very close to extinction.
5. Only the Eyebrow Wedgefish (*Rhynchobatus palpebratus*) is not assessed as Critically Endangered, due to it occurring primarily in Australia where fishing pressure is low, and some management measures are in place. Australia represents a ‘lifeboat’ for the three wedgefish and one giant guitarfish species occurring there.
6. To conserve populations and permit recovery, a suite of measures will be required which will need to include species protection, spatial management, bycatch mitigation, and harvest and international trade management, all of which will be dependent on effective enforcement.

## 1 INTRODUCTION

One of the defining features of the Anthropocene will be the loss of biodiversity, both on land and in the oceans (Dirzo et al., 2014; McCauley et al., 2015). The oceans face a wide range of threats but our understanding of how these drive population decline and extinction in individual species remains poor. There has long been concern for the extent of marine declines but relatively few local, regional, and global extinctions have been documented (Dulvy, Sadovy, & Reynolds, 2003; Dulvy, Pinnegar, & Reynolds, 2009; McCauley et al., 2015). Nevertheless, the challenges of monitoring marine species, in particular those that do not surface to breathe or do not return to land to breed (such as marine mammals, reptiles, and seabirds), may mean that marine extinctions are underestimated, and indeed humanity may be on the cusp of witnessing a marine extinction pulse (McCauley et al., 2015). Systematically evaluating extinction risk in marine species is therefore critical to understand patterns of decline and to drive management and conservation measures in an attempt to limit extinction.

The chondrichthyan fishes – sharks, rays, and ghost sharks (i.e. chimaeras) (hereafter referred to as ‘sharks and rays’) are a marine group with elevated extinction risk; an estimated quarter of species are threatened globally (Dulvy et al., 2014). This extinction risk assessment reveals that sawfishes, wedgefishes, and guitarfishes are amongst the most threatened families and are of global conservation concern (Dulvy et al., 2016; Jabado, 2018; Moore, 2017). Recent advances in taxonomy and phylogenetics have resolved some of the complex relationships of these rays (Faria et al., 2013; Last, Séret, & Naylor, 2016b; Last et al., 2016c) enabling a new assessment of their status. The order Rhinopristiformes was resurrected by Last et al. (2016b) and is now considered to consist of the sawfishes (family Pristidae), wedgefishes (Rhinidae), giant guitarfishes (Glaucostegidae), guitarfishes (Rhinobatidae), and banyo rays (Trygonorrhinidae). Collectively, these groups can be referred to as the ‘shark-like rays’ given their phylogenetic position as rays, but morphological similarities to sharks (in particular the shark-like posterior body, including dorsal and caudal fins).

An accurate assessment of extinction risk requires the delineation of taxonomic units. The sawfishes have historically been plagued by poor taxonomic resolution and species delineation (Faria et al., 2013), and similarly, the status of wedgefishes has been challenging to understand because of uncertain species identification (Jabado, 2019). The ‘whitespotted wedgefish’ (i.e. *Rhynchobatus djiddensis*) species-complex has been poorly-defined with the name ‘*Rhynchobatus djiddensis*‘ used widely for wedgefishes across the Indo-West Pacific Ocean region prior to clarification of species distributions and recognition that *R. djiddensis* is in fact restricted to the Western Indian Ocean (Last et al., 2016c). Additionally, several new wedgefish species have been recently described (Last, Ho, & Chen, 2013; Last, Kyne, & Compagno, 2016a; Séret & Naylor, 2016), and while species identification remains an issue in the field, species taxonomic boundaries and geographical distributions are now well enough defined to allow a more accurate assessment of global extinction risk.

The international trade in shark fin for the Asian soup market has incentivised targeting and retention of sharks and shark-like rays (Dent & Clarke, 2015). Sawfishes, wedgefishes, and giant guitarfishes all have ‘white’ fins, amongst the best quality and highest value in the fin trade (Dent & Clarke, 2015; Hau, Abercrombie, Ho, & Shea, 2018; Moore, 2017; Suzuki, 2002). Domestically, the meat is also an important protein source, linking the status of these species to livelihoods in developing tropical countries (Jabado, 2018; Moore, 2017; Moore, Séret, & Armstrong, 2019). Sawfishes, wedgefishes, and guitarfishes were previously common in soft-bottom habitats of shallow, warm waters, but have been heavily exploited from exposure to intensive trawl and gillnet fisheries in these habitats (Jabado, 2018; Moore, 2017).

Conservation and management measures have lagged resource exploitation in the shark-like rays. Considerable progress has recently been made in raising awareness and implementing management for sawfishes following the release of a global conservation strategy (Fordham, Jabado, Kyne, Charvet, & Dulvy, 2018; Harrison & Dulvy, 2014), and urgency has been declared for action on wedgefishes and giant guitarfishes (Moore, 2017). High levels of exploitation and the increasing pattern of targeting for the international trade has led to concern that wedgefishes and giant guitarfishes are at extinction risk levels similar to sawfishes (Hau et al., 2018; Jabado, 2018; Moore, 2017). Extinction risk assessments for sawfishes were reviewed in 2013; these highlighted rapid declines, local extinctions, and the need for serious investment in conservation and management (see Dulvy et al., 2016; Harrison & Dulvy, 2014). Extinction risk was previously assessed for most wedgefishes and giant guitarfishes between 2003 and 2007.

A global reassessment of extinction risk of all sharks and rays is being undertaken through the International Union for the Conservation of Nature (IUCN) Species Survival Commission Shark Specialist Group’s Global Shark Trends Project. Wedgefishes and giant guitarfishes were prioritised for reassessment given the issues outlined above. Here, the IUCN Red List Categories and Criteria are applied to wedgefishes and giant guitarfishes globally. First, pertinent background information (biogeography and habitat; life history; and, exploitation, use, and trade) is reviewed before summarizing population trends and IUCN Red List categories.

## 2 METHODS

### 2.1 Taxonomic scope

The taxonomic scope of this study are the 10 recognised species of wedgefishes (Rhinidae) and six giant guitarfishes (Glaucostegidae) of the order Rhinopristiformes following Last et al. (2016c) (Tables 1 & 2).

**TABLE 1.** Distribution and life history of wedgefishes (Rhinidae). Life history data from Last & Stevens (2009); Last et al. (2016c); van der Elst (1993); White & Dharmadi (2007).

| Species | Distribution | Depth range (m) | Maximum size (cm TL) | Size-at-maturity (cm TL) | Size-at-birth (cm TL) | Litter Size | Generation Length (years) |
| --- | --- | --- | --- | --- | --- | --- | --- |
| Bowmouth Guitarfish<br>*Rhina ancylostoma* Bloch & Schneider, 1801 | Indo-West Pacific | Inshore–70 | 270 | ♀ ~180<br>♂ 150–175 | 46–48 | 2–11 | 15 |
| Bottlenose Wedgefish<br>*Rhynchobatus australiae* Whitley, 1939 | Indo-West Pacific | Inshore–60 | ~300 | ♀ ~155<br>♂ 110–130 | 46–50 | 7–19<br>(mean 14) | 15 |
| Clown Wedgefish<br>*Rhynchobatus cooki* Last, Kyne & Compagno, 2016 | Southeast Asia | n/a | 81 | ♀ n/a<br>♂ <70 | n/a | n/a | 10 |
| Whitespotted Wedgefish<br>*Rhynchobatus djiddensis* (Forsskål, 1775) | Western Indian | Inshore–70 | 310 | ♀ n/a<br>♂ ~150 | 60 | 4 | 15 |
| Taiwanese Wedgefish<br>*Rhynchobatus immaculatus* Last, Ho & Chen, 2013 | Taiwan | n/a | >99† | ♀ n/a<br>♂ n/a | n/a | n/a | 10 |
| Smoothnose Wedgefish<br>*Rhynchobatus laevis* (Bloch & Schneider, 1801) | Indo-West Pacific | Inshore–60 | >200 | ♀ n/a<br>♂ ~130 | n/a | n/a | 15 |
| African Wedgefish<br>*Rhynchobatus luebberti* Ehrenbaum, 1915 | Eastern Atlantic | Inshore–35 | ~300 | ♀ n/a<br>♂ n/a | 79–85 | 2–5 | 15 |
| Eyebrow Wedgefish<br>*Rhynchobatus palpebratus* Compagno & Last, 2008 | Indo-West Pacific | 5–61 | 262 | ♀ n/a<br>♂ 103 | 46–50 | n/a | 15 |
| Broadnose Wedgefish<br>*Rhynchobatus springeri* Compagno & Last, 2010 | Southeast Asia | 16–40 | 213 | ♀ n/a<br>♂ ~115 | n/a | n/a | 15 |
| False Shark Ray<br>*Rhynchorhina mauritaniensis* Séret & Naylor, 2016 | Mauritania | n/a | 275 | ♀ n/a<br>♂ n/a | n/a | n/a | 15 |

TL, total length; n/a, not available; †Immature male, maximum size suspected to be ~150 cm TL.

**TABLE 2.** Distribution and life history of giant guitarfishes (Glaucostegidae). Life history data from Capapé & Zaouali (1994); Enajjar et al. (2012); Gohar & Mazhar (1964); Last et al. (2016c); Moore et al. (2012); Moore & Peirce (2013); Muhammad Moazzam Khan, pers. comm., 07/02/2019; Prasad (1951); Seck et al. (2004).

| Species | Distribution | Depth range (m) | Maximum Size (cm TL) | Size-at-maturity (cm TL) | Size-at-birth (cm TL) | Litter Size | Generation Length (yrs) |
| --- | --- | --- | --- | --- | --- | --- | --- |
| Blackchin Guitarfish<br>*Glaucostegus cemiculus* (Geoffroy St Hilaire, 1817) | Eastern Atlantic & Mediterranean Sea | Inshore–80 | 265 | ♀ 163 (Senegal)<br>♀ 110–138 (Tunisia)<br>♂ 155 (Senegal)<br>♂ 100–112 (Tunisia) | ~34 | 16–24 (Senegal)<br>5–12 (Tunisia) | 15 |
| Sharpnose Guitarfish<br>*Glaucostegus granulatus* (Cuvier, 1829) | Northern Indian | Inshore–120 | 229 | ♀ n/a<br>♂ n/a | ~39 | 6–18 | 15 |
| Halavi Guitarfish<br>*Glaucostegus halavi* (Forsskål, 1775) | Northern Indian | Inshore–100 | 187 | ♀ ~83<br>♂ ~83 | ~29 | up to 10 | 10 |
| Widenose Guitarfish<br>*Glaucostegus obtusus* (Müller & Henle, 1841) | Indo-West Pacific | Inshore–60 | 93 | ♀ n/a<br>♂ ~48 | n/a | 4–10 | 10 |
| Clubnose Guitarfish<br>*Glaucostegus thouin* (Anonymous, 1798) | Indo-West Pacific | Inshore–60 | ~300 | ♀ n/a<br>♂ n/a | n/a | n/a | 15 |
| Giant Guitarfish<br>*Glaucostegus typus* (Bennett, 1830) | Indo-West Pacific | Inshore–100 | 270 | ♀ 150–180<br>♂ 150–180 | 38–40 | n/a | 15 |

TL, total length; n/a, not available

### 2.2 Application of the IUCN Red List Categories and Criteria

The IUCN Red List Categories and Criteria (Version 3.1) were applied following the Guidelines for Using the IUCN Red List Categories and Criteria (IUCN, 2012; IUCN Standards and Petitions Subcommittee, 2017). Assessments were undertaken at the global level, i.e. for the entire global population of each species. For each species, data on taxonomy, distribution, population status, habitat and ecology, major threats, use and trade, and conservation measures were collated from the peer-reviewed literature, fisheries statistics, grey literature, and consultation with species and fisheries experts.

Draft assessments were prepared in the IUCN Species Information Service (SIS) online database. Each assessment was peer-reviewed by at least two reviewers who were trained in the application of the IUCN Red List Categories and Criteria and who were familiar with shark-like rays and the fisheries interacting with them. A summary of the assessments was also provided to the entire IUCN Species Survival Commission Shark Specialist Group (SSG) for their consultation and input (174 members). Assessments were then submitted to the IUCN Red List Unit (Cambridge, UK) where they underwent further review and quality checks before being accepted for publication on the IUCN Red List (version 2019-2, July 2019, www.iucnredlist.org; IUCN, 2019).

The IUCN Red List applies eight extinction risk categories: Extinct (EX), Extinct in the Wild (EW), Critically Endangered (CR), Endangered (EN), Vulnerable (VU), Near Threatened (NT), Least Concern (LC), and Data Deficient (DD) (IUCN, 2012; Mace et al., 2008). A species is considered EX ‘when there is no reasonable doubt that the last individual has died’; EW ‘when it is known only to survive in cultivation, in captivity or as a naturalised population (or populations) well outside the past range’; CR, EN, and VU species are considered to be facing an extremely high, very high, or high risk of extinction in the wild, respectively; NT species do ‘not qualify for CR, EN or VU now, but is close to qualifying for or is likely to qualify for a threatened category in the near future’; LC species do not qualify for CR, EN, VU, or NT; finally, DD species have ‘inadequate information to make a direct, or indirect, assessment of its risk of extinction based on its distribution and/or population status (IUCN, 2012).

Each species was assessed against the five Red List criteria: A – population size reduction; B – geographic range size; C – small population size and decline; D – very small or restricted population; and, E – quantitative analysis (for example, population viability analysis) (see IUCN, 2012; IUCN Standards and Petitions Subcommittee, 2017; Mace et al., 2008). To qualify for one of the three threatened categories (CR, EN, or VU), a species has to meet a quantitative threshold for that category in any of the five criteria listed above (A–E). A collation and review of available information indicated that there were no data available to assess species under criteria C, D, or E, and these criteria are therefore not considered further here. All species were assessed under criterion A, with some consideration of criterion B for range restricted species.

Criterion A applies a set of quantitative thresholds to consider population reduction scaled over a period of three generation lengths (3 GL) (IUCN Standards and Petitions Subcommittee, 2017; Mace et al., 2008). While there are a range of demographic approaches to calculating generation length (IUCN Standards and Petitions Subcommittee, 2017), these are generally data intensive and have not been applied to any wedgefish or giant guitarfish. Therefore, to derive generation length (GL), a simple measure that requires only female age-at-maturity and maximum age was used:

GL = ((maximum age – age-at-maturity)/2)) + age-at-maturity

This value represents the median age of parents of the current cohort. To derive population reduction over 3 GL, the proportional decline over the *x* years of available catch rate or landings datasets was calculated and this was used to calculate annual proportional change, which was then scaled across the 3 GL period.

### 2.3 Distribution mapping

A global distribution map (Appendix I) was generated for each species, primarily following the ranges in Last et al. (2016c), with some minor modifications based on new records. Ranges were clipped to the maximum depth of each species, and for those wedgefishes without known depth ranges, these were set to the maximum confirmed depth of the family (70 m; Table 1). To determine global patterns of biodiversity, species richness maps were produced for all species combined, wedgefishes only, and giant guitarfishes only. All maps were prepared using ArcMap 10.4 (ESRI, 2016).

### 2.4 Calculation of a Red List Index

A Red List Index (RLI) was calculated based on the number of species in each Red List category at each of three time periods. The index was calculated as the weighted sum of species status scaled by the number of species. An ‘equal-step’ weighting was used where the weight (*W_c_*) equals zero for LC, 1 - NT, 2 - VU, 3 - EN, 4 - CR, and 5 - EX or EW. Hence, a species moving from LC to NT will contribute as much to the index as a species moving from EN to CR. The RLI is scaled to range from 1 (where all species are LC) to 0 (where all species are EX), and is calculated as:

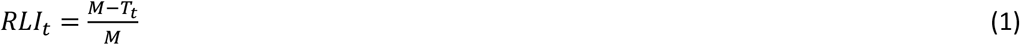

where *M* is the maximum threat score, which is the number species multiplied by the maximum weight assigned to EX species (here, a value of 5), and in this case for 16 species is 16 × 5 = 80. The current threat score (*T_t_*) is the sum of the number of species in each threat category in year *t* (*N_c(t)_*), times the category weight (*W_c_*).

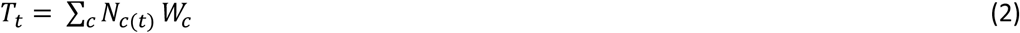

Hence, the threat score for the current assessment would be calculated as the *N_c(t)_* = 15 species that are Critically Endangered (*W_c_* = 4), giving 4 × 15 = 60, summed with the one Near Threatened species (*W_c_* = 1). Thus, the current threat score *T_t=2019_* is 60 + 1 = 61 and the *RLI*_*t*=2019_ = (80 − 61) / 80 = 0.2375.

Retrospective assessments were developed for two earlier time periods, which were chosen as 2005 and 1980 (with the current assessments set at 2020). Prior to this current reassessment, all six giant guitarfishes and seven of the wedgefishes had assessments published on the IUCN Red List (wedgefishes: 1 EN, 6 VU; giant guitarfishes: 1 EN, 5 VU). All changes in Red List category were considered to be non-genuine changes as a result of new information (IUCN Standards and Petitions Subcommittee, 2017). In other words, if what is currently understood was known during the previous assessments, it is likely that the assigned status of those species would have been different. ‘Back casting’ is undertaken by retrospectively assigning status based on current understanding of the spatial and temporal pattern of coastal human population growth, the development of general fishing pressure, an understanding of the availability of fishing gear capable of capturing sharks and rays, and the development of the international trade demand for shark and shark-like ray fins (e.g. Blaber et al., 2019; Clarke, Milner-Gulland, & Bjorndal, 2007; Cripps, Harris, Humber, Harding, & Thomas, 2015; Sousa, Marshall, & Smale, 1997; Stewart et al., 2010).

Red List Indices were also calculated for the two main oceanic regions, the Indo-West Pacific Ocean region (hereafter, ‘Indo-West Pacific’), and the Eastern Atlantic Ocean and Mediterranean Sea region (hereafter, ‘Eastern Atlantic’), as well as individually for each of the 87 countries containing some proportion of at least one of the 16 species assessed here. Threat scores applied to the two oceanic regions followed the equal-step weighting outlined above. For disaggregating the global RLI to the national level, the equation is amended such that:

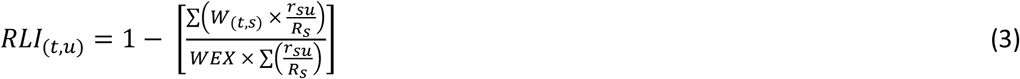

where *t* is the year of assessment *u*, is the country and *W*_(*t*,*s*)_ is the Red List threat at year *t* for each species, multiplied by 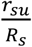, which represents the proportion of each species’ total range found within the Exclusive Economic Zone (EEZ) of each country. This is summed across all species found in each country’s EEZ and divided by the maximum threat score (*WEX* = 5), multiplied by the sum of proportional species’ ranges. The final RLI value is derived from subtracting by 1 so that higher RLI values indicate less negative changes in Red List status across species and vice versa (as with the global RLI). Finally, the national conservation responsibility for all species were calculated separately for each of the two oceanic regions, based on the sum of all threat scores across species within a country multiplied by each of the species’ proportional ranges for that country. Resulting national responsibility values were normalized to range between 0 and 1 for both regions.

## 3 RESULTS

Here, summaries of (1) biogeography and habitat; (2) life history; (3) exploitation, use and trade; (4) population status; (5) IUCN Red List Categories; (6) the possible extinction of two wedgefish species; and, (7) the Red List Index for wedgefishes and giant guitarfishes, are presented.

### 3.1 Biogeography and habitat

The Indo-West Pacific is the centre of diversity for wedgefishes (8 species) and giant guitarfishes (5 species), with the remaining three species occurring in the Eastern Atlantic (including the Mediterranean Sea for the Blackchin Guitarfish (*Glaucostegus cemiculus*)) (Tables 1 & 2, Figure 1, Appendix I). Distributions range from extremely widespread, i.e. Bowmouth Guitarfish (*Rhina ancylostoma*) and Bottlenose Wedgefish (*Rhynchobatus australiae*) to the very restricted, i.e. Taiwanese Wedgefish (*Rhynchobatus immaculatus*), Clown Wedgefish (*Rhynchobatus cooki*), and False Shark Ray (*Rhynchorhina mauritaniensis*) (Appendix I). These latter three species are known only from fish landing sites in northern Taiwan, Singapore and Jakarta, and Mauritania, respectively (Last et al., 2013; 2016a; Séret & Naylor, 2016), and therefore their exact distributions remain undefined. *Rhynchorhina mauritaniensis* is potentially the most range-restricted species, as it is currently only known from a single location, the Banc d’Arguin National Park in Mauritania (Séret & Naylor, 2016).

**FIGURE 1.**
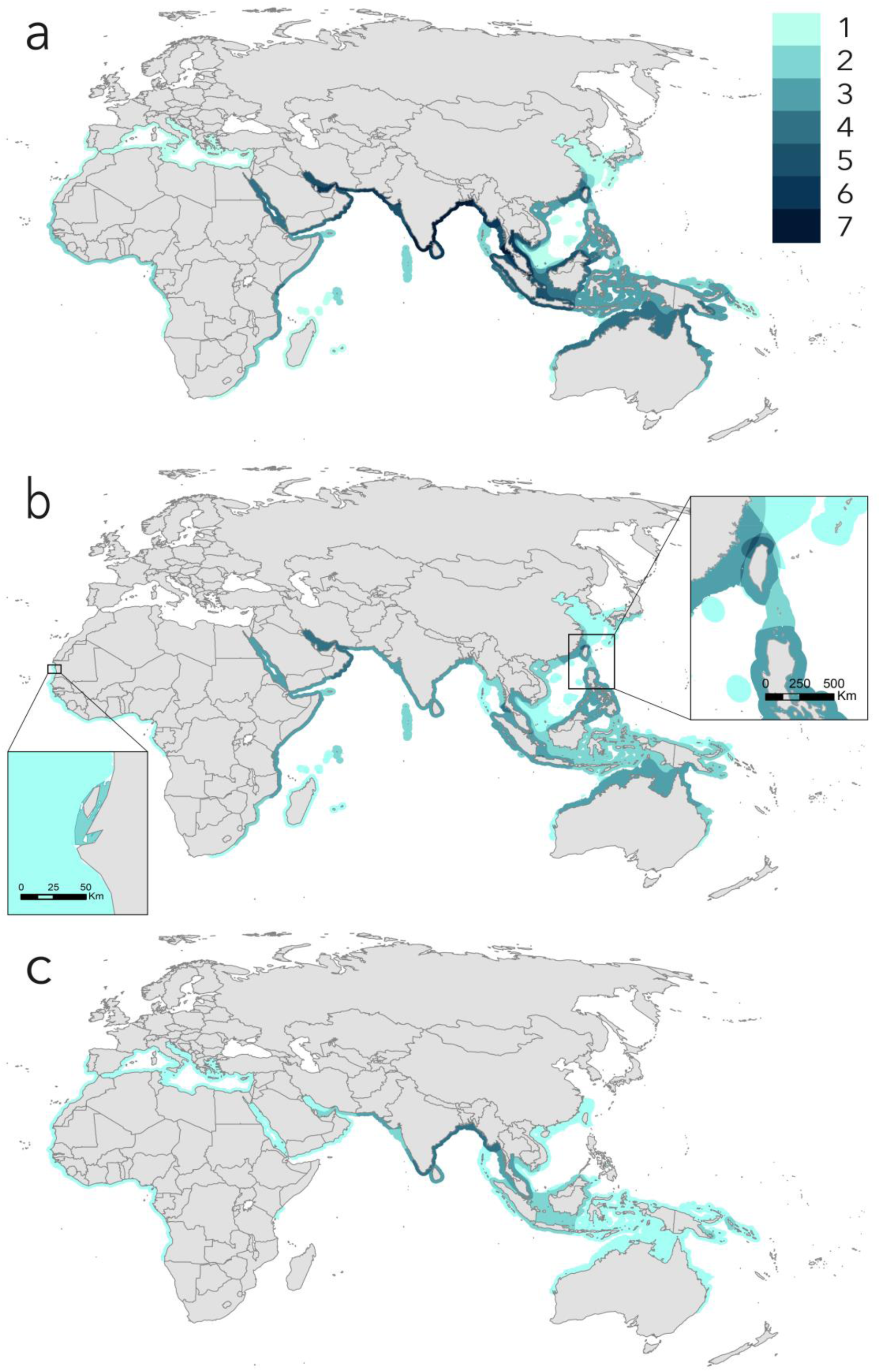
Wedgefish and giant guitarfish species richness: (a) Global species richness of wedgefishes and giant guitarfishes combined (n = 16 species); (b) Global species richness of wedgefishes (n = 10 species); (c) Global species richness of giant guitarfishes (n = 6 species).

Both families primarily occur in tropical to warm temperate waters from close inshore to the mid continental shelf, although two species (*R. ancylostoma, R. australiae*) are also known to occur around island chains far from continental landmasses; wedgefishes occur to a maximum depth of at least 70 m (although exact depth ranges are unknown for three species) and giant guitarfishes to a maximum of 120 m (Tables 1 & 2; Last et al., 2016c). Some species have been recorded from the estuarine reaches of rivers and the Broadnose Wedgefish (*Rhynchobatus springeri*) is thought to be a habitat specialist of shallow brackish coastal and estuarine waters (Compagno & Last, 2010), while others can be associated with coral reefs (e.g. *R. ancylostoma*).

### 3.2 Life history

The life history of wedgefishes and giant guitarfishes is generally very poorly known, with only a limited number of dedicated studies on aspects of their biology and ecology, with the exception of *G. cemiculus*. Wedgefishes are large species, with most species reaching >200 cm total length (TL) and up to 310 cm TL in the Whitespotted Wedgefish (*Rhynchobatus djiddensis*), although *R. cooki* is an exceptionally small species (81 cm TL), while maximum size is unknown for *R. immaculatus* (the largest collected specimen was still immature at 99 cm TL) (Table 1). Giant guitarfishes reach 300 cm TL (Clubnose Guitarfish, *Glaucostegus thouin*) with most species >200 cm TL, except the Halavi Guitarfish (*Glaucostegus halavi*; 187 cm TL) and the Widenose Guitarfish (*Glaucostegus obtusus*; 93 cm TL). Size-at-maturity and size-at-birth are poorly-known with data gaps for most species (Tables 1 & 2).

Reproduction is lecithotrophic viviparous in both families with generally small, but variable litter sizes: in the wedgefishes, from as low as 2 pups per litter in *R. ancylostoma* (range: 2–11) and the African Wedgefish (*Rhynchobatus luebberti*) (2–5), to as high as 19 pups per litter in *R. australiae* (7–19), and in the giant guitarfishes, from a low of 4 pups per litter in *G. obtusus* (4–10) to as high as 24 pups per litter in *G. cemiculus* (Tables 1 & 2). *Glaucostegus cemiculus* exhibits some regional variation with 16– 24 pups per litter in Senegal and 5–12 in Tunisia. Litter sizes are available for only 4 of 10 wedgefishes and 4 of 6 giant guitarfishes. Reproductive periodicity is suspected to be annual in *G. cemiculus* (Capapé & Zaouali, 1994), but periodicity, and therefore annual fecundity, are largely unknown across the two families.

There is a general lack of age and growth data. For wedgefishes, the only study (White, Simpfendorfer, Tobin, & Heupel, 2014) was based on mixed samples of *R. australiae* and the Eyebrow Wedgefish (*Rhynchobatus palpebratus*), and therefore has limited biological meaning. Maximum observed age was 12 years (female of 183 cm TL) (White et al., 2014) which would be well below longevity given that *R. palpebratus* reaches 262 cm TL and *R. australiae* reaches ∼300 cm TL. For giant guitarfishes, a maximum observed age for the Giant Guitarfish (*Glaucostegus typus*) of 19 years (250 cm TL female) was reported by White et al. (2014) and while age-at-maturity was not reported, it can be estimated from the growth curve as 7 years (by reading the corresponding age at 165 cm TL, the mid-point of size-at-maturity; Last et al., 2016c). This estimate is not sex-specific as the growth curve of White et al. (2014) was based on combined sexes. For *G. cemiculus*, a maximum observed age of 15 years (198 cm TL female), a male of age-at-maturity of 2.9 years, and a female age-at-maturity of 5.1 years was reported by Enajjar, Bradai, & Bouain (2012).

An estimate of generation length (GL) for *G. cemiculus* of 9.5 years based on the age data of Enajjar et al. (2012) is likely an underestimate given maximum observed age was for an individual well below maximum size (198 vs 265 cm TL). This GL estimate does, however give a suitable estimate for smaller (<200 cm TL) wedgefish and giant guitarfish species. A GL estimate for *G. typus* of 13 years based on the age data of White et al. (2014) is a reasonable estimate given that maximum observed age was for an individual close to maximum size (250 vs. 270 cm TL) (White et al., 2014). To ensure consistency across IUCN Red List Assessments, 15 years was applied as an estimated GL to large (≥200 cm TL) species, and 10 years for smaller species (<200 cm TL).

### 3.3 Exploitation, use, and trade

Globally, wedgefishes and giant guitarfishes are subject to intense fishing pressure on their coastal and shelf habitats (Stewart et al., 2010) that is unregulated across the majority of their distributions. They are captured in industrial, artisanal, and subsistence fisheries with multiple fishing gears, including gillnet, trawl, hook and line, trap, and seine net and are generally retained for their meat and fins (Bonfil & Abdallah, 2004; Jabado, 2018; Moore, 2017). There is a high level of fisheries resource use and increasing fishing pressure which has resulted in the over-exploitation or depletion of demersal coastal fisheries resources in significant areas of the Indo-West Pacific and the Eastern Atlantic, including West Africa, India, and Southeast Asia (FAO, 2018b; Mohamed & Veena, 2016; Pauly & Chuenpagdee, 2003; Stewart et al., 2010; Stobutzki et al., 2006). The major exception is Australia where fishing pressure is considerably lower (this is also the case for some smaller range states such as New Caledonia, and South Africa which are at the geographic limit of the range of a small number of species).

In general, fishing effort and the number of fishers has increased in recent decades across the range of these species, with demand for shark and ray products increasing over the same period due to the shark fin trade (Chen, 1996; Diop & Dossa, 2011; Jabado et al., 2017). Several examples of this increase from across the global range of wedgefishes and giant guitarfishes include (1) Mauritania which has seen a significant increase in fishing effort since the second half of the 20th Century: in 1950 there were 125 pirogues (small-scale fishing boats), rising to nearly 4,000 in 2005 (Belhabib et al., 2012); (2) Senegal, where the number of artisanal pirogues rose from ∼5,000 in 1982 to 12,699 in 2006, although it has since fallen slightly to 11,889 in 2013 (ANSD, 2016; FAO, 2008); (3) Madagascar, where the number of pirogues rose from ∼5,000 in 1983 to ∼22,000 in 1996 (Cooke, 1997); (4) the Red Sea, where the number of traditional boats tripled from 3,100 to 10,000 from 1988 to 2006 (Bruckner, Alnazry, & Faisal, 2011); and, (5) the Indian state of Gujarat, where the number of trawlers increased from about 6,600 in the early 2000s to 11,582 in 2010 (CMFRI, 2010; Jabado et al., 2017; Zynudheen, Ninan, Sen, & Badonia, 2004). This increasing fishing effort has put significant pressure on all species. Furthermore, the high value of fins is driving retention and trade of wedgefishes and giant guitarfishes globally, with these species targeted in the Mediterranean Sea, West Africa, East Africa, India, and the Indo-Malay Archipelago, among other places (Barrowclift, Temple, Stead, Jiddawi, & Berggren, 2017; Diop & Dossa, 2011; IOTC, 2005; Jabado, 2018; Lteif, 2015; Moore, 2017; Newell, 2016; Seisay, 2005).

Both the meat and fins drive utilisation and trade. The high-quality meat is consumed by many coastal communities in tropical countries and it is also dried, salted, and consumed locally or traded internationally (e.g. Moore, 2017; Jabado, 2018). Large whole wedgefishes (>200 cm total length; TL) have been traded for a high value of up to US$680 each (e.g. Jabado, 2018). Prices for the highly-valued ‘white’ fins of large shark-like rays are reportedly as high as US$964/kg (Jabado, 2019). Other reported prices include US$396/kg for wedgefish fins (Chen, 1996) and an average price of US$276/kg and US$185/kg for *Qun chi* (fins from shark-like rays) in Guangzhou (mainland China) and Hong Kong, respectively (Hau et al., 2018). In addition to meat and fins, other uses include the skin which may be dried and traded internationally as a luxury leather product (Haque, Biswas, & Latifa, 2018), the eggs which are sometimes dried and consumed locally, the heads which may be dried and used as either fish meal or fertilizer (Haque et al., 2018; R.W. Jabado, unpubl. data), and the snout of giant guitarfishes are considered a delicacy in Singapore where they are steamed, and the gelatinous filling consumed.

### 3.4 Population status

#### 3.4.1 Data availability

Where rhinopristoid rays have been targeted or exploited as incidental catch, severe declines, population depletions, and localised disappearances have occurred (e.g. Dulvy et al., 2016; Jabado, 2018; Moore, 2017; Tous, Ducrocq, Bucal, & Feron, 1998). However, there are no species-specific time-series data available that can be used to calculate population reduction in wedgefishes and giant guitarfishes. Despite this, there are a number of relevant historical accounts and contemporary datasets for landings and catch rates. All of these accounts and datasets are from the Indo-West Pacific (from Iran to Indonesia), but can also be considered informative for understanding population reduction in wedgefishes and giant guitarfishes more broadly where they are under heavy exploitation, including in the Eastern Atlantic. The five contemporary datasets are available for landings data or catch rates at varying levels of taxonomic resolution (e.g. ‘guitarfishes’, ‘whitespotted wedgefishes’ etc.) from Iran, Pakistan, western and eastern India, and Indonesia. These datasets likely include various species of wedgefishes and giant guitarfishes and in each case, probable species are listed below. One dataset (Raje & Zacharia, 2009) does not include rhinopristoids but rather presents landings data for myliobatoid rays (stingrays, eagle rays, butterfly rays, and devil rays). However, this can be used to infer declines in wedgefishes and giant guitarfishes given overlapping distributions, habitat, and susceptibility to capture in the same fishing gear. A summary of these datasets and corresponding proportional decline over 3 GL is provided in Table 3.

**TABLE 3.** Overall decline, annual proportional change, proportion remaining, and proportional decline over three generation lengths for landings and catch rate datasets. Proportional decline is provided for small (<200 cm TL) and large (≥200 cm TL) wedgefish and giant guitarfish species by applying a 3 generation length of 30 and 45 years, respectively.

| Location | Iran |  | Sindh, Pakistan |  | Balochistan, Pakistan |  | Maharashtra, India |  | Indonesia |  |
| --- | --- | --- | --- | --- | --- | --- | --- | --- | --- | --- |
| Data type | Landings (t) |  | Landings (t) |  | Landings (t) |  | Catch rate (kg/hr) |  | Landings (t) |  |
| Data category | 'giant guitarfish' |  | 'rhinobatid' |  | 'rhinobatid' |  | 'myliobatoid rays' |  | 'whitespotted wedgefishes' |  |
| Data period (x years) | 1997–2016 |  | 1999–2011 |  | 1994–2011 |  | 1990–2004 |  | 2005–2015 |  |
| Data source | FAO (2018a) |  | M.A. Gore, unpubl. data |  | M.A. Gore, unpubl. data |  | Raje & Zacharia (2009) |  | DGCF (2015; 2017) |  |
| Proportional decline over x years | 0.665 |  | 0.720 |  | 0.806 |  | 0.631 |  | 0.876 |  |
| Annual proportional change | 0.947 |  | 0.907 |  | 0.913 |  | 0.936 |  | 0.827 |  |
| 3 generation lengths (3GL) | 30 yrs | 45 yrs | 30 yrs | 45 yrs | 30 yrs | 45 yrs | 30 yrs | 45 yrs | 30 yrs | 45 yrs |
| Proportion remaining | 0.194 | 0.086 | 0.053 | 0.012 | 0.065 | 0.016 | 0.136 | 0.050 | 0.003 | 0.0002 |
| Proportional decline over 3GL | 0.806 | 0.914 | 0.947 | 0.988 | 0.935 | 0.984 | 0.864 | 0.950 | 0.997 | 0.9998 |

#### 3.4.2 Indo-West Pacific

##### 3.4.2.1 Historical accounts

Research trawl survey data from the Gulf of Thailand showed a 93% decline in catch rates of ‘Rhinobathidae’ (a name that is likely to include wedgefishes and guitarfishes broadly) over a short time period from peak catches in 1968 to a low in 1972 (Pauly, 1979, Ritragsa, 1976). Similarly, catch rates of ‘rays’ declined by 92% from 1963 to 1972. Secondly, the Indonesian Aru Islands wedgefish gillnet fishery rapidly expanded from its beginnings in the mid-1970s to reach its peak in 1987 with more than 500 boats operating before catches then declined very rapidly leading to only 100 boats left fishing in this area in 1996 (Chen, 1996). In all likelihood, the fleet redistributed to other areas as wedgefishes were depleted and catch rates declined. Thirdly, investors in Indonesia withdrew from a wedgefish fishery in the Malaku and Arafura Seas because the resource had been overfished by 1992 resulting in limited returns for their investment (Suzuki, 2002). Lastly, research trawl surveys in the Java Sea showed the decline of ‘rays’ between 1976 and 1997 by ‘at least an order of magnitude’ (i.e., a decline of at least 90%) (Blaber et al., 2009). It is worth noting that recent trawl surveys in the Java Sea recorded only a single individual *Rhynchobatus* (Tirtadanu, Suprapto, & Suwarso, 2018), and in the North Natuna Sea (north of the Java Sea), trawl surveys recorded only two individuals (Yusup, Priatna, & Wagiyo, 2018).

##### 3.4.2.2 Iran landings dataset

Landings data for the ‘giant guitarfish’ category are available from Iran for 1997–2016 (20 years; FAO, 2018a; Table 3). This grouping likely includes all rhinids (wedgefishes) and glaucostegids (giant guitarfishes) occurring locally, including *R. ancylostoma*, *R. australiae*, *R. djiddensis*, Smoothnose Wedgefish (*Rhynchobatus laevis*), Sharpnose Guitarfish (*Glaucostegus granulatus*), and *G. halavi*. Landings declined by 67% over this period, the equivalent of an 81% and 91% population reduction over the last 3 GL of smaller species (30 years) and larger species (45 years), respectively.

##### 3.4.2.3 Pakistan landings dataset

Landings data for the ‘rhinobatid’ category are available from Pakistan for 1993–2011 (19 years) covering the country’s two coastal provinces (data collated from Pakistan Government records; M.A. Gore, unpubl. data; Table 3). This grouping likely includes all rhinids and glaucostegids occurring locally, including *R. ancylostoma*, *R. australiae*, *R. laevis*, *G. granulatus*, *G. halavi*, and *G. obtusus*, as well as rhinobatids (guitarfishes) including Bengal Guitarfish (*Rhinobatos annandalei*). Data from Sindh province showed a 72% decrease from peak landings in 1999 to a low in 2011, the equivalent of a 95% and 99% population reduction over the last 3 GL of smaller species (30 years) and larger species (45 years), respectively. Data from Balochistan province showed an 81% decrease from peak landings in 1994 to the last data point in 2011, the equivalent of a 94% and 98% population reduction over the last 3 GL of smaller species (30 years) and larger species (45 years), respectively. No wedgefish or giant guitarfish were observed during surveys of southwest Balochistan fish landings between 2007 and 2010 (M.A. Gore and U. Waqas, unpubl. data).

##### 3.4.2.4 Western India ray catch rate dataset

Catch rate data for myliobatoid rays (this includes a variety of demersal rays, but does not include rhinopristoids) are available from Maharashtra, western India for 1990–2004 (15 years; Raje and Zacharia, 2009; Table 3). The catch rate declined by 63% over this period (despite fishing effort doubling during this time), the equivalent of an 86% and 95% population reduction over the last 3 GL of smaller species (30 years) and larger species (45 years), respectively.

##### 3.4.2.5 Eastern India landings dataset

Landings data for ‘guitarfishes’ are available from Tamil Nadu, eastern India for 2002–2006 (5 years; Mohanraj, Rajapackiam, Mohan, Batcha, & Gomathy, 2009). This grouping was reported in the paper to include *R. ancylostoma*, ‘*R. djiddensis*’ (which would therefore include *R. australiae* and *R. laevis*, since *R. djiddensis* does not occur in this area), *G. granulatus*, and *G. obtusus*, but was also likely to include *G. thouin* and *G. typus*. Landings declined by 86% over this period. Furthermore, species-specific trawl landings data were reported for ‘*R. djiddensis*’ (i.e. *R. australiae* and *R. laevis*), with a decline of 87% over this period. This time-period is however too short to derive an equivalent population reduction over three generations.

##### 3.4.2.6 Indonesia landings dataset

Landings data for ‘whitespotted wedgefishes’ are available from Indonesia for 2005–2015 (11 years; DGCF, 2015; 2017; Table 3). This grouping likely includes *R. ancylostoma*, *R. australiae*, *R. cooki*, *R. palpebratus*, and *R. springeri*. It may also include giant guitarfishes, but in any case, the trends can be considered representative of giant guitarfishes occurring locally due to overlapping habitat and catchability (i.e. *G. obtusus*, *G. thouin*, and *G. typus*). Landings declined by 88% over this period, the equivalent of >99% population reduction over the last 3 GL of both smaller species (30 years) and larger species (45 years). An additional data point available for 2016 is excluded from this analysis. This datum suggests a massive increase in reported landings which is an artefact of the inclusion of a wider range of rays in the reported figure (DGCF, 2017; Muhammad Anas, pers. comm., 11/2/2019).

##### 3.4.2.7 East Africa anecdotal reports

The above information spans Iran to Southeast Asia, with less information available from East Africa in the Western Indian Ocean. Anecdotal reports from this region suggest that artisanal longline fishing led to declines in *R. djiddensis* in southern Mozambique (which was one of the main target species of the fishery) as this species was abundant on reefs before longline fisheries began in the early 2000s and subsequently, are only seen in low numbers (Pierce et al., 2008). In Zanzibar, fisher interviews indicated that there were perceived declines in wedgefish or that they are rare (Schaeffer, 2004); wedgefishes were a retained bycatch of commercial prawn trawling in Tanzania (Rose, 1996). Intense fishing pressure across the Tanzanian shelf has likely resulted in population reduction, mirroring those outlined above for the Indo-West Pacific more broadly. In Madagascar, there was a decrease in the size of wedgefish caught in artisanal fisheries over time (Humber et al., 2017), though this could be due, in part, to the targeting of larger individuals. A steep decline in catch-per-unit-effort (CPUE) can be inferred from reported catch reductions from 10–20 sharks per day in 1992 to 1–3 sharks per day in 1995 in Morondava, West Madagascar, with fishers subsequently moving further afield to fish (Cooke, 1997). Wedgefish, a high-value target species, would likely have declined by a similar order of magnitude as sharks. In South Africa, there was a marked decline in CPUE of *R. djiddensis* in shark bather protection nets in KwaZulu-Natal during the period 1979–2017 (Nomfundo Nakabi, pers. comm., 17/04/2018). This decline is not considered to be a good indicator of population reduction as it may be explained, at least partially, by a shift in gear deployment whereby nets were gradually lifted off the substrate (which would reduce the capture of demersal species).

##### 3.4.2.8 Australia

The one region in which wedgefish and giant guitarfish populations may be in a better state than most of the rest of their range is Australia. Here, fishing effort is relatively low, the use of turtle exclusion devices in trawl fisheries reduces the catch of large rays (Brewer et al. (2006) recorded a reduction of 94%), and there are some controls on wedgefish catch and retention. Estimates of fishing mortality rates for wedgefish and giant guitarfish species in the Northern Prawn Fishery (the largest Australian fishery to interact these species) are well below those that would lead to significant population declines (Zhou & Griffiths, 2008).

#### 3.4.3 Eastern Atlantic and Mediterranean Sea

Data on population status in the Eastern Atlantic Ocean and Mediterranean Sea is sparse, but there are several lines of evidence to support similar population reductions, as well as local extinctions. In the Mediterranean Sea, *G. cemiculus* was regarded as historically common within both northern (de Buen, 1935; Doderlein, 1884) and southern (Bradaï, Saidi, Enajjar, & Bouain, 2006; Quignard & Capapé, 1971; Whitehead, Bauchot, Hureau, Nielsen, & Tortonese, 1984) areas. However, there are now contrasting situations between these two areas. The species has largely disappeared from the northern Mediterranean Sea and was not recorded in extensive trawl surveys under the Mediterranean International Trawl Surveys (MEDITS) program from 1994 to 2015 (Newell, 2016; Relini & Piccinetti, 1991), nor in trawl surveys in the Adriatic Sea between 1948 and 2005 (Ferretti, Osio, Jenkins, Rosenberg, & Lotze, 2013). In the southern Mediterranean Sea (including the Gulf of Gabés and areas of the eastern Mediterranean, which seem to be core parts of the species’ distribution), the species is still present and, in some areas, still commonly caught (e.g. Echwikhi, Saidi, & Bradaï, 2014; Lteif, 2015; Newell, 2016; Soldo, Briand, & Rassoulzadegan, 2014).

In West Africa, trend data are lacking, but evidence points to severe declines of wedgefishes*. Rhynchobatus luebberti* is known to have disappeared from a significant part of West Africa (Mauritania to Sierra Leone but apparently with the exception of the Banc d’Arguin National Park; Diop & Dossa, 2011). However, the species is now sparsely reported in the Banc d’Arguin National Park with only two individuals recorded in the past decade during fish landing site monitoring (the most recent record being February 2019) (Sall Amadou, pers. comm., 14/02/19; Saïkou Oumar Kidé, pers. comm., 14/02/19). This species was moderately abundant across its former range in the 1960s but declined thereafter (Bernard Séret, pers. comm., 07/02/19); during Guinean trawl surveys in the 1960s, catch rates were as high as 30–34 kg/hr (William, 1968). By contrast, recent fish market surveys across the region have either failed to locate it or found only low numbers of individuals. In The Gambia, annual surveys from 2010 to 2018 of landing sites that regularly land guitarfishes and other rays have not recorded the species (Moore et al., 2019). In one artisanal demersal gillnet fishery in Mayumba, Gabon (between 30 to 40 boats), surveys between February 2013 and October 2015 identified 40 individuals, and surveys between May and October 2018 identified 5 individuals (Godefroy de Bruyne, pers. comm., 14/09/18). Observers on board national trawlers off Gabon have not recorded the species in monitoring which commenced in 2015, despite many species of rays being recorded (Emmanuel Chartrain, pers. comm., 15/02/19). In Port Gentil, Gabon (around 400 boats), where rays are targeted, *R. luebberti* has not been seen during ongoing surveys that commenced in June 2017 (Godefroy de Bruyne, pers. comm., 14/09/18). A 2006 capture by a recreational fishing guide in Guinea-Bissau was reportedly described as ‘very, very rare’ (Moore, 2017). It was also recently confirmed from Sao Tomé Island through a photographic record (Reiner & Wirst, 2016).

### 3.5 IUCN Red List categories

All wedgefishes and giant guitarfishes were assessed as CR A2, with the exception of *R. palpebratus* which was assessed as NT (nearly meeting criterion A2). That is, 15 out of 16 species are inferred to have undergone a population reduction of >80% over the last three generations (30–45 years), where ‘the causes of reduction may not have ceased OR may not be understood OR may not be reversible’ (IUCN, 2012). In this case, the causes are understood (over-exploitation in target and bycatch fisheries, driven by human consumption and trade in meat and fins), they are theoretically reversible (through the implementation of management measures; see Discussion), but they have not ceased (largely unregulated exploitation continues with fishing effort increasing). These population reductions are based on ‘an index of abundance appropriate to the taxon’ (IUCN, 2012), i.e. the declines in landings and catch rates presented above, and ‘actual or potential levels of exploitation’ (IUCN, 2012), i.e. high levels of exploitation in target and bycatch fisheries. Red List categories and criteria along with a brief assessment justification for wedgefishes are provided in Table 4 and for giant guitarfishes in Table 5.

**TABLE 4.** Summary of IUCN Red List (RL) Categories and Criteria for wedgefishes (Rhinidae).

| Species | RL Category & Criteria | Justification |
| --- | --- | --- |
| *Rhina ancylostoma* | CR A2bd | Indo-West Pacific population reductions in rhinopristsoids; high levels of exploitation across most of range; some refuge in Australia but not considered a large enough proportion of range to lower assessment |
| *Rhynchobatus australiae* | CR A2bd | Indo-West Pacific population reductions in rhinopristsoids; high levels of exploitation across most of range; some refuge in Australia but not considered a large enough proportion of range to lower assessment |
| *Rhynchobatus cooki* | CR A2bd | Indo-West Pacific population reductions in rhinopristsoids; high levels of exploitation across range; only a single record since 1996 in well surveyed and heavily fished areas |
| *Rhynchobatus djiddensis* | CR A2bd | Indo-West Pacific population reductions in rhinopristsoids; high levels of exploitation across most of range; some refuge in South Africa but not considered a large enough proportion of range to lower assessment |
| *Rhynchobatus immaculatus* | CR A2bd | Indo-West Pacific population reductions in rhinopristsoids; limited distribution in heavily fished area; no refuge |
| *Rhynchobatus laevis* | CR A2bd | Indo-West Pacific population reductions in rhinopristsoids; high levels of exploitation across most of range; no refuge |
| *Rhynchobatus luebberti* | CR A2d | Once common and now only sporadically recorded; localised extinction; high levels of exploitation across range; no refuge |
| *Rhynchobatus palpebratus* | NT A2bd | Assuming disjunct range of Australia/PNG, Thailand & Taiwan (as opposed to wider Australasian/Southeast Asian range): high levels of exploitation in Thailand & Taiwan, refuge in northern Australia (significant proportion of range) |
| *Rhynchobatus springeri* | CR A2bd | Indo-West Pacific population reductions in rhinopristsoids; high levels of exploitation across range; no refuge |
| *Rhynchorhina mauritaniensis* | CR A2d | High levels of exploitation across range; absence of records; no refuge |

CR, Critically Endangered; NT, Near Threatened.

**TABLE 5.** Summary of IUCN Red List (RL) Categories and Criteria for giant guitarfishes (Glaucostegidae).

| Species | RL Category & Criteria | Justification |
| --- | --- | --- |
| *Glaucostegus cemiculus* | CR A2d | Localised extinctions in northern Mediterranean; high levels of exploitation across West Africa; no refuge |
| *Glaucostegus granulatus* | CR A2bd | Indo-West Pacific population reductions in rhinopristsoids; high levels of exploitation across most of range; no refuge |
| *Glaucostegus halavi* | CR A2bd | Indo-West Pacific population reductions in rhinopristsoids; high levels of exploitation across most of range |
| *Glaucostegus obtusus* | CR A2bd | Indo-West Pacific population reductions in rhinopristsoids; high levels of exploitation across range; no refuge |
| *Glaucostegus thouin* | CR A2bd | Indo-West Pacific population reductions in rhinopristsoids; high levels of exploitation across range; rarity; no refuge |
| *Glaucostegus typus* | CR A2bd | Indo-West Pacific population reductions in rhinopristsoids; high levels of exploitation across most of range; some refuge in Australia but not considered a large enough proportion of range to lower assessment |

CR, Critically Endangered.

Parts of Australasia and South Africa stand apart as the clear exceptions to the widespread intense fisheries elsewhere. Four species (*R. ancylostoma*, *R. australiae*, *R. palpebratus*, and *G. typus*) occur in tropical and warm-temperate waters of Australia where fishing pressure is relatively low and fisheries management measures are in place. For widely-distributed species (*R. ancylostoma*, *R. australiae*, and *G. typus*) this proportion of the species’ range is not considered to be large enough relative to the global range to lower the global CR assessment status. The bulk of the currently recognised distribution of *R. palpebratus* is within Australian waters, influencing its more favourable global status of NT, compared to the other species. It should be noted however, that the full distribution of this species is not well understood, and the disjunct records (Australia/New Guinea, Thai Andaman Sea, and Taiwan; Compagno & Last, 2008; Ebert et al., 2013; Last et al., 2016c) suggests that it is/was more widely ranging throughout Southeast Asia and Australasia, or that there is an unresolved taxonomic issue. Fishing pressure is high where *R. palpebratus* occurs outside of Australia and based on the landings and catch rate data presented above, it is inferred that the species has undergone a >80% population reduction over the last three generations (45 years) in the Asian part of its range. There is little contemporary information on the species outside of Australia, and it has not been recorded in recent landing site surveys on the Andaman coast of Thailand (Shin Arunrugstichai, pers. comm., 16/01/19). If the species was in fact wider-ranging throughout the Indo-Malay Archipelago/Southeast Asia, as its disjunct distribution suggests, it would likely have undergone a population reduction over the last three generations high enough to qualify it for a threatened category (possibly as high as CR, the status of all other wedgefishes).

Generally, there are few catch and trend data for elasmobranchs in the Eastern Atlantic and there was no population trend information available for the three species found there: *R. luebberti*, *R. mauritaniensis*, and *G. cemiculus*. Nevertheless, inference can be drawn from general regional fisheries trends. Fishing effort and the number of fishers has increased in recent decades across West Africa, with demand for shark and ray product increasing over the same period due to the shark fin trade (Diop & Dossa, 2011). For example, large regional fishing nations including Mauritania and Senegal have seen significant increases in fishing effort since the second half of the 20th Century, with considerable artisanal and industrial fishing fleets operating in waters off West Africa (ANSD, 2016; Belhabib et al., 2012; FAO, 2008; ONS, 2017). The severe population reductions inferred for Indo-West Pacific wedgefishes and giant guitarfishes from several datasets could likely be considered representative of the situation in the Eastern Atlantic. Indeed, heavy exploitation has led to the depletion of *R. luebberti* and the possible disappearance of *R. mauritaniensis*.

### 3.6 Possible extinction of two wedgefish species

The most at-risk species are those with very-restricted ranges: *R. cooki* of the Indo-Malay Archipelago and *R. mauritaniensis* of Mauritania, both of which may be very close to extinction, even before they were taxonomically described in 2016 (Last et al., 2016a; Séret & Naylor, 2016). The full distribution of *R. cooki* is unclear as it has only been collected from fish landing sites in Singapore and Jakarta (Indonesia), and these landings come from fisheries that operate widely across the Indo-Malay Archipelago (Last et al., 2016a). There has only been a single record of this species since 1996 (an individual observed at a Singapore fish market in early 2019; Naomi Clark-Shen and Kathy Xu, pers. comm., 26/05/2019). The limited number of records in a heavily fished and scientifically well-surveyed area raises serious concerns for the species. Further surveys are required to understand its contemporary occurrence and status, and ongoing monitoring of fish markets should pay special attention to wedgefish landings while making an effort to determine from fishers where the species was caught, and therefore its natural range.

*Rhynchorhina mauritaniensis* is known only from one location, the Banc d’Arguin National Park, Mauritania. The Indigenous Imraguen population of the local area were traditionally subsistence fishers until a shift to commercial shark fishing from the mid-1980s (see Belhabib et al., 2012; Diop & Dossa, 2011). This shift, along with increasing artisanal and industrial fishing effort in Mauritanian waters possibly depleted the population even before it was formally described by Séret & Naylor (2016). This species is known to occur in an area where targeting of sharks has been prohibited since 2003 (Diop & Dossa, 2011) and only Indigenous fishers are permitted to fish using traditional methods (the Banc d’Arguin National Park). However, the artisanal fishing effort in the National Park, combined with illegal fishing effort is considerable (Belhabib et al., 2012), and *R. mauritaniensis* is known to be landed locally. Individuals have been observed with their fins removed when landed, and the fins sold to local fin dealers (Séret & Naylor, 2016). This species is not likely to have any refuge from fishing within its very restricted range given the combined effort from subsistence, artisanal, and illegal fishing coupled with the high value of its fins. The species’ extent of occurrence is estimated to be <5,000 km^2^, which combined with its presence in only one location, and an inferred continuing decline in the number of mature individuals due to this ongoing fishing pressure, meets EN under criterion B (as EN B1ab(v)) (IUCN, 2012). However, a lack of records, high actual levels of exploitation, and a broad understanding of declines of similar species in the Indo-West Pacific, as well as the locally-occurring *R. luebberti*, also lead us to infer that *R. mauritaniensis* has undergone a >80% population reduction over the last three generations (45 years) and is assessed as CR A2d.

The poorly-known *R. immaculatus* is also considered to be at elevated risk. It is another species known only from fishing landing sites, in this case, in northern Taiwan (Last et al., 2013). The lack of records suggests a very limited distribution which raises serious concerns for its ability to sustain historic and current levels of fishing pressure. Taiwan is a major fishing nation with a long history of exploitation of coastal resources, which were considered to be overfished by the 1950s (and which led to the development of Taiwan’s distant water fleet) (Kuo & Booth, 2011). Taiwan ranks among the top 20 shark fishing nations globally (Lack & Sant, 2011) and is a major global shark fin trading nation (Clarke et al., 2006; Dulvy et al., 2014). Furthermore, there is an extensive illegal, unreported, and unregulated (IUU) fishing issue in Taiwan (Kuo & Booth, 2011).

### 3.7 Red List Index

The global RLI for wedgefishes and giant guitarfishes starts relatively high in 1980 at 0.7, declining steadily to 0.43 in 2005 and further to 0.24 in the current assessment (2020) (Figure 3a). The global index is driven mainly by the greater diversity of the Indo-West Pacific, which has a similar RLI in 1980 of 0.63. In the Eastern Atlantic however, a steep decline in RLI occurs between 1980 to 2005, from 1 to 0.4, compared to the Indo-West Pacific, which declines from 0.63 to 0.43 over the same time period (Figure 3a). This difference in decline rates is likely due to the later development of wedgefish and giant guitarfish fisheries and fin trade in the Eastern Atlantic. By 1980, it is inferred that 11 species were already likely to be threatened (i.e. Red List category of CR, EN, or VU); all these species occur in the Indo-West Pacific, where there has been an early development of fisheries and trade, particularly in Asia with its proximity to Hong Kong as the major shark fin trade centre. For example, *R. immaculatus* (Indo-West Pacific), is inferred as already CR by 1980 due to the early development of intensive fisheries in Taiwan and proximity to Hong Kong. By contrast, all three species found in the Eastern Atlantic were LC in 1980 (thus resulting in RLI of 1 for the region; Figure 3a). By 2005, it was inferred that at a global level, one species was CR, 13 were EN, one was VU, and one was NT. By the current assessment (2020), the RLI has declined to 0.25 and 0.2 for the Indo-West Pacific, and the Eastern Atlantic, respectively (Figure 3a).

**FIGURE 2.**
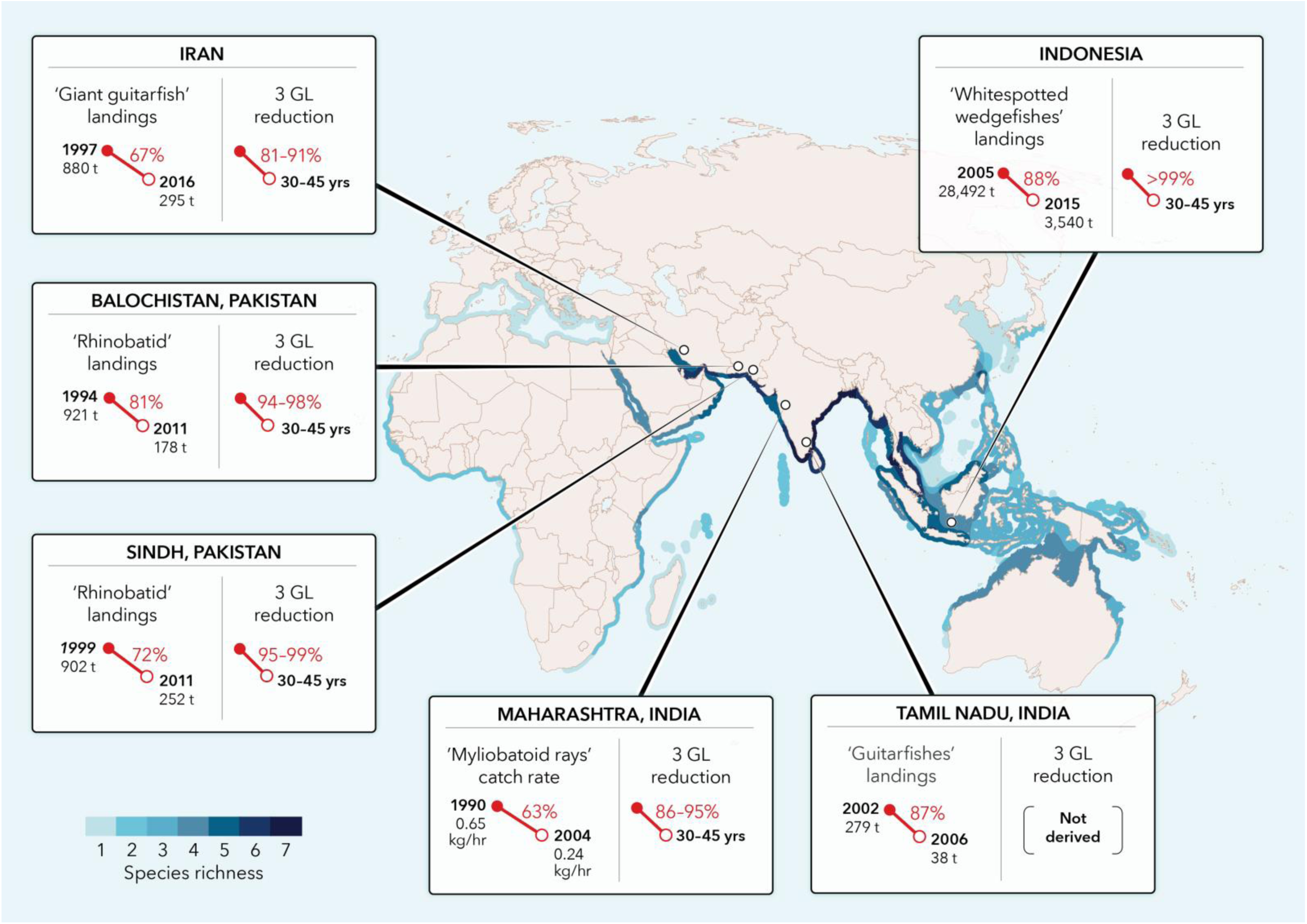
Summary of landings and catch rate data used to infer population reductions in wedgefishes and giant guitarfishes overlaid on the map of global species richness of wedgefishes and giant guitarfishes combined (Figure 1A). Data sources are provided in Table 3. GL, generation length.

**FIGURE 3.**
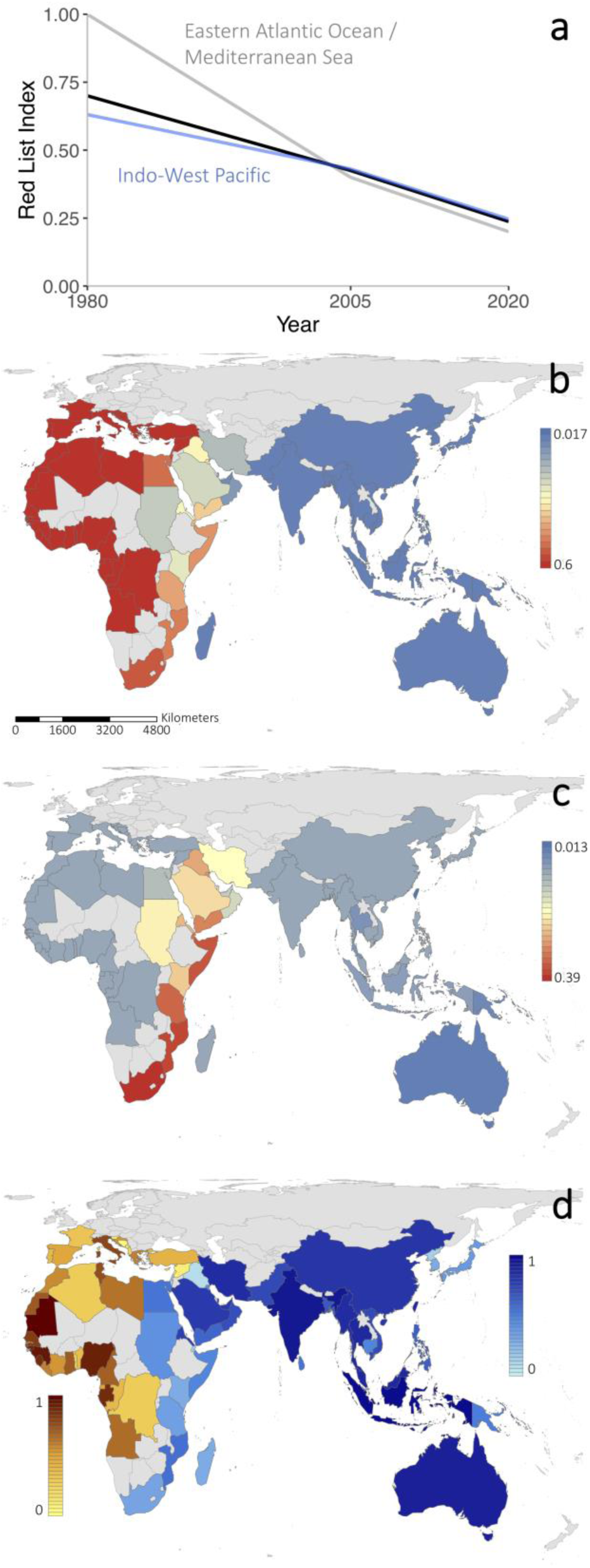
Red List Indices for wedgefishes and giant guitarfishes. (a) Global Red List Index (RLI; black line) decomposed for the two main oceanic regions, Indo-West Pacific Ocean (blue line), and the Eastern Atlantic Ocean and Mediterranean Sea (gray line); decline in country-weighted RLI from (b) 1980 to 2005, and (c) 2005 to 2020; and, (d) National conservation responsibilities for all wedgefish and giant guitarfish species across the two main regions, Indo-West Pacific Ocean (blues) and the Eastern Atlantic Ocean and Mediterranean Sea (yellows).

The trends in wedgefish and giant guitarfish fisheries and fin trade described above are reflected in the geographic regions that display the sharpest declines in Red List Index between the different assessment years (Figure 3b and c). Declines in RLI between 1980 and 2005 are concentrated in West African and Mediterranean Sea nations (Figure 3b), shifting to East African nations between 2005 and 2020 (Figure 3c). Declines are less severe in the Indo-West Pacific since most species are already likely threatened by 1980 (Figure 3b and c). Species’ ranges in the Eastern Atlantic and the Indo-West Pacific overlap with the EEZ of forty-one and forty-six nations, respectively (Figure 3d, Table 6). The top ten percent of nations in the Eastern Atlantic responsible for the conservation of species in this region are Mauritania, Guinea, Guinea-Bissau, and Nigeria (collectively representing 57% of all national responsibility for the region); in the Indo-West Pacific, these nations are Indonesia, India, Australia, Taiwan, and Malaysia (representing 55% of all responsibility for the region; Figure 3d, Table 6).

**TABLE 6.** National conservation responsibilities (NCR) for all wedgefish and giant guitarfish species across 87 countries. Conservation responsibility (calculated as the sum of threat scores for each species weighted by the proportion of species range contained within each country’s EEZ; see methods) is determined separately and normalised to range from 0 to 1 for comparability between the two distinct regions, the Eastern Atlantic Ocean and Mediterranean Sea region, and Indo-West Pacific Ocean region.

| Eastern Atlantic and Mediterranean Sea |  | Indo-West Pacific |  |
| --- | --- | --- | --- |
| Country | NCR | Country | NCR |
| Mauritania | 1 | Indonesia | 1 |
| Guinea | 0.209 | India | 0.487 |
| Guinea-Bissau | 0.172 | Australia | 0.322 |
| Nigeria | 0.149 | Taiwan, Province of China | 0.242 |
| Gabon | 0.112 | Malaysia | 0.221 |
| Sierra Leone | 0.108 | Thailand | 0.202 |
| Egypt | 0.101 | Myanmar | 0.177 |
| Senegal | 0.086 | Islamic Republic of Iran | 0.166 |
| Italy | 0.067 | China | 0.155 |
| Tunisia | 0.066 | Saudi Arabia | 0.136 |
| Ghana | 0.063 | United Arab Emirates | 0.122 |
| Western Sahara | 0.062 | Eritrea | 0.086 |
| Cameroon | 0.050 | Pakistan | 0.082 |
| Angola | 0.048 | Vietnam | 0.080 |
| Libya | 0.044 | Oman | 0.073 |
| Greece | 0.041 | Bangladesh | 0.069 |
| Liberia | 0.040 | Qatar | 0.068 |
| Morocco | 0.036 | Philippines | 0.064 |
| Croatia | 0.031 | Yemen | 0.056 |
| Côte d'Ivoire | 0.030 | Mozambique | 0.048 |
| Equatorial Guinea | 0.028 | Egypt | 0.040 |
| Spain | 0.027 | Sri Lanka | 0.037 |
| Gambia | 0.023 | Papua New Guinea | 0.030 |
| Turkey | 0.019 | Somalia | 0.022 |
| Congo | 0.016 | Cambodia | 0.022 |
| Portugal | 0.015 | Kuwait | 0.020 |
| Benin | 0.009 | Sudan | 0.018 |
| France | 0.008 | Japan | 0.016 |
| Algeria | 0.007 | Bahrain | 0.015 |
| The Democratic Republic of the Congo | 0.007 | United Republic of Tanzania | 0.011 |
| Togo | 0.005 | South Africa | 0.009 |
| Israel | 0.004 | Kenya | 0.007 |
| Albania | 0.004 | Republic of Korea | 0.007 |
| Cyprus | 0.002 | Madagascar | 0.006 |
| Montenegro | 0.002 | Seychelles | 0.004 |
| Lebanon | 0.001 | Brunei Darussalam | 0.003 |
| Syrian Arab Republic | 0.001 | Maldives | 0.002 |
| Malta | 0.001 | Singapore | 0.002 |
| Slovenia | <0.001 | Israel | 0.002 |
| Bosnia and Herzegovina | <0.001 | Democratic People's Republic of Korea | 0.002 |
| Monaco | 0 | Djibouti | 0.001 |
|  |  | Solomon Islands | 0.001 |
|  |  | Iraq | 0.001 |
|  |  | Timor-Leste | <0.001 |
|  |  | Mauritius | <0.001 |
|  |  | Réunion | 0 |

## 4 DISCUSSION

This study brings together several lines of evidence to show severe population reductions in wedgefishes and giant guitarfishes globally, resulting in 15 of 16 species (94%) facing an ‘extremely high risk of extinction’, i.e. assessed as Critically Endangered on the IUCN Red List. That makes these the most imperilled marine fish families globally, overtaking the sawfishes which are comprised of three CR and 2 EN species (IUCN, 2019). The demand for shark and ray products, including the high-value ‘white’ fins of wedgefishes and giant guitarfishes will continue to drive and incentivise targeting and retention, and urgent action is required to prevent extinctions. Next, the following topics are considered: (1) data quality and knowledge gap issues in assessing extinction risk in wedgefishes and giant guitarfishes; (2) the intersection between species richness and threat; (3) the current shortfall in conservation and management; (4) Australia as a refuge for a quarter of the fauna; and, 1. (5) measures that are needed to prevent extinction.

### 4.1 Data quality and knowledge gaps

Most of the available data upon which these assessments were based were catch landings under broad aggregate categories such as ‘giant guitarfish’, ‘rhinobatid’, and ‘whitespotted wedgefishes’. These non-species-specific groupings limit the possibility of analysing population trends for individual species but are useful to infer trends based on overlapping habitat and depth ranges across species, and likely similar catchability in extensive coastal and shelf fisheries in tropical and warm temperate Indo-West Pacific and Eastern Atlantic waters.

Although landings data are not a direct measure of abundance, these can be used to infer population reduction where landings have decreased while fishing effort has remained stable or increased, hence approximating a decline in CPUE. In nearly all cases used here to assess population status, there was no reason to suspect that overall effort had decreased (although directed fishing effort may have shifted in response to resource collapse/depletion; e.g. the Aru Islands gillnet fishery in Indonesia). In fact, fishing effort and power is continuing to increase globally as the coastal human population continues to grow and fishing technology and market access improves. Some of the highest increases in fishing effort and power occur in the Asian region (Anticamara, Watson, Gelchu, & Pauly, 2011; Watson et al., 2013), which is a centre of wedgefish and giant guitarfish diversity. Hence, declining catches are inferred to likely indicate reductions in abundance.

All of the wedgefishes and giant guitarfishes were assessed using the IUCN Red List ‘Population size reduction’ A criterion (IUCN, 2012; IUCN, 2019; Mace et al., 2008). The IUCN Red List Criteria were designed to allow a range of data quality to be used, allowing taxa to be assessed in the absence of complete, high-quality datasets (IUCN Standards and Petitions Subcommittee, 2017). Moving from the highest to the lowest levels of acceptable data quality, IUCN accepts information that is ‘observed’ (e.g. population decline based on well-documented observations of all known individuals in the population); ‘estimated’ (e.g. population decline based on repeated surveys that involve statistical assumptions); ‘projected’ (e.g. a future population decline model based on past repeated surveys and threats that are unlikely to stop); ‘inferred’ (e.g. a population decline based on trade or fisheries landings data), or ‘suspected’ (e.g. information based on circumstantial evidence). For the wedgefishes and giant guitarfishes, population reductions were ‘inferred’. Of the available contemporary datasets, only the catch rate data of myliobatoid rays from Maharashtra, India (Raje & Zacharia 2009) could be used to ‘estimate’ a population reduction (86–95% over three generations). However, when applied to the assessment of wedgefish and giant guitarfish extinction risk, the data quality was low since population reductions were inferred from another demersal ray lineage (Myliobatiformes). Because the datasets used from Iran, Pakistan, and Indonesia (DGCF, 2015; 2017; FAO, 2018a; M.A. Gore, unpubl. data) consisted of landings only, these could only be used to ‘infer’ population reduction.

Inferring population reductions from broad landings data of aggregate species categories highlighted the data deficiency around these species, not only in catch and trade data, but also in basic habitat and life history parameters. For example, amongst the wedgefishes, depth ranges are completely unknown for three species, annual fecundity is unknown across the family (and litter size is known from only four species), and generation length had to be inferred from giant guitarfishes. Across both families, age and growth studies are restricted to only two published works (Enajjar et al., 2012; White et al., 2014), with no accurate data for wedgefishes given that White et al. (2014) analysed mixed species samples.

### 4.2 The intersection between species richness and threat

Species richness is highest in areas of significant fishing effort, and these hotspots of overlap between diversity and pressure may be priorities for management. The Indo-West Pacific (13 species) is the centre of diversity for wedgefishes and giant guitarfishes, with low diversity in the Eastern Atlantic (three species), and no species in the Western Atlantic or Eastern Pacific. The Northern Indian Ocean, particularly the Arabian/Persian Gulf to India, and the Indo-Malay Archipelago are areas of special concern. These regions include several countries that rank among the top 20 shark fishing nations globally, specifically Indonesia, India, Pakistan, Malaysia, Thailand, Sri Lanka, and Iran (Lack & Sant, 2011) and are under high levels of coastal fishing effort (Stewart et al., 2010). Unsurprisingly, there have been steep declines in shark and ray landings over the past decade in this region likely due to the collapse of chondrichthyan stocks (Davidson, Krawchuk, & Dulvy, 2016) It is informative to consider the sheer number of fishing vessels in operation in these regions, for example (1) all Indian states have high numbers of fishing vessels (e.g. as reported in 2010: Maharashtra, 5,613 trawlers; Kerala, 3,678 trawlers, Tamil Nadu, 5,767 trawlers; total trawlers in India: 35,228) and a high number of gillnetters (total of 20,257 as reported in 2010), (2) Oman with 19,000 artisanal boats, (3) Pakistan with 2,000 trawlers, (4) Sri Lanka with 24,600 gillnet vessels operating in 2004; and, (5) Indonesia with ∼600,000 fishing vessels in marine waters (CMFRI, 2010; Dissanayake, 2005; Jabado et al., 2017; KKP, 2016). The intensity of fishing pressure on the coastal and shelf waters leaves little refuge for wedgefishes and giant guitarfishes.

While fishing pressure is the primary threat driving population reduction of wedgefishes and giant guitarfishes, these effects are compounded by habitat loss and degradation. The shallow, inshore soft-bottom habitat preferred by the species is threatened by habitat loss and environmental degradation (Jabado et al., 2017; Moore, 2017; Moore, McCarthy, Carvalho, & Peirce, 2012; Stobutzki et al., 2006; White & Sommerville, 2010). In the Arabian Sea and adjacent waters, dredging and coastal land reclamation has increased in recent years and has resulted in almost total loss of mangroves in some areas, such as Bahrain (Jabado et al., 2017; Sheppard et al., 2010), while Southeast Asia has seen an estimated 30% reduction in mangrove area since 1980 (FAO, 2007; Polidoro et al., 2010). Combined with targeted and bycatch fishing, the cumulative impacts of habitat loss and degradation will hinder recovery.

### 4.3 Current shortfall in conservation and management

There are minimal international and national management measures in place for wedgefishes and giant guitarfishes, and these are not at the scale currently required to curtail the severe extinction risk of these species. Regarding international agreements, *R. australiae* was listed on Appendix II of the Convention on the Conservation of Migratory Species of Wild Animals (CMS) in 2017 which aims to provide a framework for the coordination of measures adopted by Range States to improve the conservation of the species. However, listing is not the same as implementation; a recent review of implementation of CMS listings revealed serious deficiencies in implementation across Range States (Lawson & Fordham, 2018). The CMS Memorandum of Understanding on the Conservation of Migratory Sharks also lists *R. australiae*, *R. djiddensis*, and *R. laevis* on Annex 1 (since December 2018). Annex 1 lists species that have an unfavourable conservation status and would significantly benefit from collaborative international conservation action. *Glaucostegus cemiculus* is listed on Annex II of the Specially Protected Areas and Biological Diversity Protocol for the Mediterranean under the Barcelona Convention, and cannot be retained on board, trans-shipped, landed, transferred, stored, sold, displayed or offered for sale, and must be released unharmed and alive (to the extent possible). European Union (EU) vessels are prohibited from fishing for guitarfishes in EU waters of several International Council for the Exploration of the Sea (ICES) sub-areas.

At the national or subnational level, there are very limited species-specific conservation or management measures in place. Some localised protections, trawl bans, finning bans, as well as general fisheries management and marine protected areas likely benefit these species, although in many areas, effective enforcement is an ongoing issue. Of 87 countries whose waters are home to one or more species of wedgefish or giant guitarfish, only eight have specific national or subnational level protections in place: (1) Guinea, where *R. luebberti* is protected (specified within the annual national fisheries management plan rather than the Fisheries Code); (2) South Africa, where *R. djiddensis* is protected; (3) Israel, where all sharks and rays are protected; (4) the United Arab Emirates (UAE), where all wedgefishes and guitarfishes are protected; (5) Kuwait, where all rays are protected; (6) Pakistan, where all guitarfishes and wedgefishes are protected in Balochistan province, and where juvenile guitarfishes and wedgefishes (less than 30 cm) are protected in Sindh province (note that this size limit is below the known size-at-birth of all wedgefishes and most giant guitarfishes; Tables 1 & 2); (7) India, where ‘*R. djiddensis*’ is protected; and, (8) Bangladesh, where ‘*R. djiddensis*’ and *G. granulatus* are protected. However, *R. djiddensis* does not occur in India or Bangladesh (Last et al., 2016c), and the species present there, *R. australiae* and *R. laevis*, are currently not listed on national legislation. Collectively, these countries represent only 19% of all conservation responsibility in the Indo-West Pacific and just 8% in the Eastern Atlantic (Israel and Guinea only).

The UAE, Qatar, and Oman have banned trawling in their waters, Malaysia has banned trawling in inshore waters, and other countries have seasonal trawl closures that may benefit species. Finning (i.e. removing fins and discarding the body at sea) has been banned in several range states including some West African countries, UAE, Oman, Iran, Israel, and Australia. This may have reduced the retention of animals solely for their fins, but fins are still traded when whole animals are landed. Furthermore, unreported finning of sharks and ‘guitar sharks’ has been reported in the Mauritania industrial shrimp fishery (Goudswaard & Meissa, 2006) and no doubt occurs more widely.

### 4.4 Lifeboat Australia

Across the global range of wedgefishes and giant guitarfishes, Australia offers some refuge for the four species occurring there (*R. ancylostoma*, *R. australiae*, *R. palpebratus*, and *G. typus*), particularly as Australia has the third highest conservation responsibility for all species occurring in the Indo-West Pacific. Fishing pressure is considerably lower in the tropical and subtropical waters of the northern half of the Australian continent than most places in the Indo-West Pacific, although the degree of connectivity with Indonesia and elsewhere is unknown. If animals regularly move into Indonesian waters, they would face significantly higher levels of fishing pressure. There are no target fisheries for these species in Australia, although they are taken as bycatch in numerous non-target fisheries (e.g. Stobutzki, Miller, Heales, & Brewer, 2002; White, Heupel, Simpfendorfer, & Tobin, 2013). The introduction of turtle exclusion devices in northern and eastern Australian prawn trawl fisheries is likely to have significantly reduced the mortality of these species in trawl fishing gear (Brewer et al., 2006). Furthermore, in the state of Queensland there is a trip limit of five wedgefishes in commercial net fisheries (DAFF, 2009) and in all jurisdictions, there are prohibitions on retention of any shark product in several fisheries. General recreational shark and ray possession limits are also in place. Lastly, Australia has a system of marine protected areas stretching across the distribution of wedgefishes and *G. typus*, and although these are multi-use parks, they include areas with limitations on fishing activities. Collectively, this management seascape may offer these species a ‘lifeboat’, a term first used by Fordham et al. (2018) in the context of Australia and sawfishes.

### 4.5 Preventing extinction

The application of IUCN Red List Categories and Criteria to wedgefishes and giant guitarfishes has shown that without immediate action, there is an extremely high likelihood of global extinction for most species. Declines in Red List Indices are severe at global, regional, and national levels, with a relatively small number of countries responsible for the majority of conservation of these species. Accurate extinction risk assessments are essential to inform policy and decision making, and to improve conservation efforts and sustainable management of shark-like rays. It is therefore necessary to continue to refine future assessments by resolving taxonomic issues, improving our understanding of species distributions and life histories, and monitoring threats.

Taxonomic resolution combined with accurate species-specific identification would greatly enhance gathering life history and habitat data, and lead to improved fisheries monitoring data recording. However, accurate identification is wanting, particularly in the ‘whitespotted wedgefish’ species-complex. While *R. ancylostoma* and *R. mauritaniensis* are distinctive, the eight *Rhynchobatus* species are morphologically similar externally, and are usually separated, if at all, by the patterning of spots around a black pectoral marking. The problem with separating these species based on spot patterns is that these may change with growth and natural variations between animals. Further compounding the matter is the poor original descriptions for many of these species; two *Rhynchobatus* species (*R. djiddensis, R. laevis*) were described over 215 years ago, and two others (*R. luebberti, R. australiae*) were described 114 and 80 years ago, respectively. In the past 11 years four new species (*R. cooki, R. immaculatus, R. palpebratus, R. springeri*) have been described, but most were based on smaller juvenile specimens, without consideration of ontogenetic changes in spot patterning. The giant guitarfishes are even more problematic since all were described more than 175 years ago, with their descriptions being poor. A taxonomic revision of both families is needed with corresponding field identification guides to improve specific-species data collection.

International trade in highly prized and valuable fins is a major driver of over-exploitation in wedgefishes and giant guitarfishes (Dent & Clarke, 2015; Hau et al., 2018; Jabado, 2018; 2019; Moore, 2017; Suzuki, 2002) and hence, trade regulation is an important part of the solution to reduce incentives to serially deplete populations of these species. Two species of wedgefish (*R. australiae* and *R. djiddensis*) and two species of giant guitarfish (*G. cemiculus* and *G. granulatus*) have been proposed for listing under Appendix II of the Convention on the International Trade in Endangered Species, with all other members of both families to be listed under the ‘look alike’ criterion. An Appendix II listing enables international trade to be controlled through export permits issued by Parties where ‘the specimen was legally obtained and if the export will not be detrimental to the survival of the species’ (CITES, 2019). There are currently 183 Parties to CITES so this instrument has broad global reach (CITES, 2019), yet implementation and enforcement are ongoing issues.

A logical first step to guide and prioritise actions for these species is a global conservation planning exercise. A global sawfish strategy was instrumental in catalysing research and monitoring for sawfishes (Fordham et al., 2018; Harrison & Dulvy, 2014), although much work remains to be done to secure those species. To conserve wedgefish and giant guitarfish populations and to permit recovery, a suite of national, regional, and international measures will be required which will need to include species protection, spatial management, bycatch mitigation, and harvest and international trade management measures. Effective enforcement of measures will require ongoing training and capacity-building (including improving species identification; Jabado, 2019). Catch monitoring, especially in artisanal fisheries, is needed to help understand local population trends and inform management. The dire situation of two wedgefish species, *R. cooki* and *R. mauritaniensis*, outlined here highlights the urgency of global concerted action.

## ACKNOWLEDGEMENTS

This work was funded by the Shark Conservation Fund as part of the Global Shark Trends Project. We thank all members of the IUCN Species Survival Commission Shark Specialist Group and other experts who contributed to the data collation to inform Red List assessments. In particular, we extend our gratitude to Sall Amadou, Shin Arunrugstichai, Emmanuel Chartrain, Naomi Clark-Shen, Godefroy de Bruyne, Katie Gledhill, Adrian Gutteridge, Saïkou Oumar Kidé, Peter Last, Muhammad Moazzam Khan, Alec Moore, Nomfundo Nakabi, Bernard Séret, Umer Waqas, and Kathy Xu. PMK was supported by the Marine Biodiversity Hub, a collaborative partnership supported through funding from the Australian Government’s National Environmental Science Program. MAG was supported by a Darwin Initiative grant. DAE was supported by the South African Institute for Aquatic Biodiversity and the Save Our Seas Foundation. NKD was supported by Natural Science and Engineering Research Council Discovery and Accelerator Awards, the Canada Research Chairs Program, and the Disney Conservation Fund.

---

**APPENDIX I.**
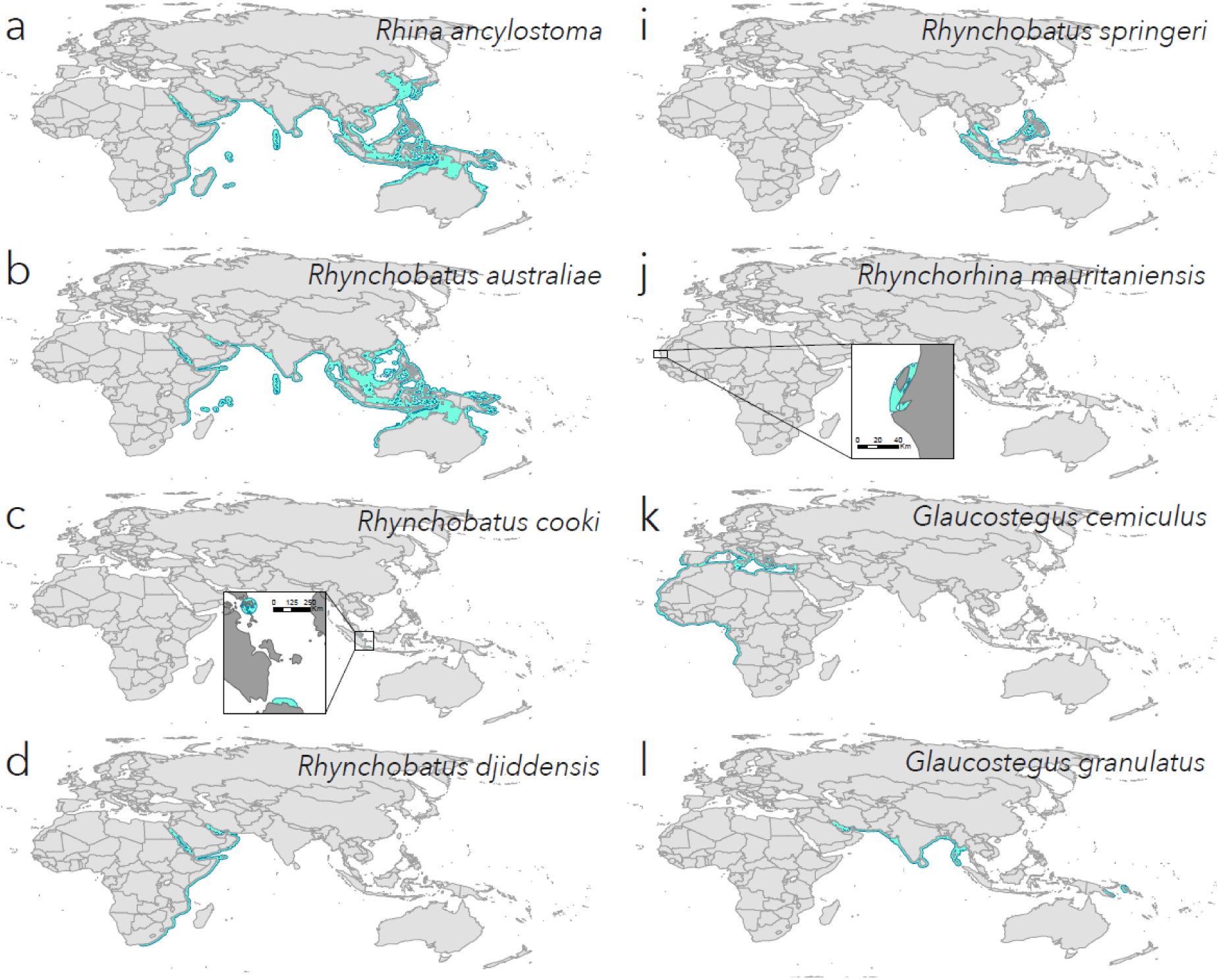
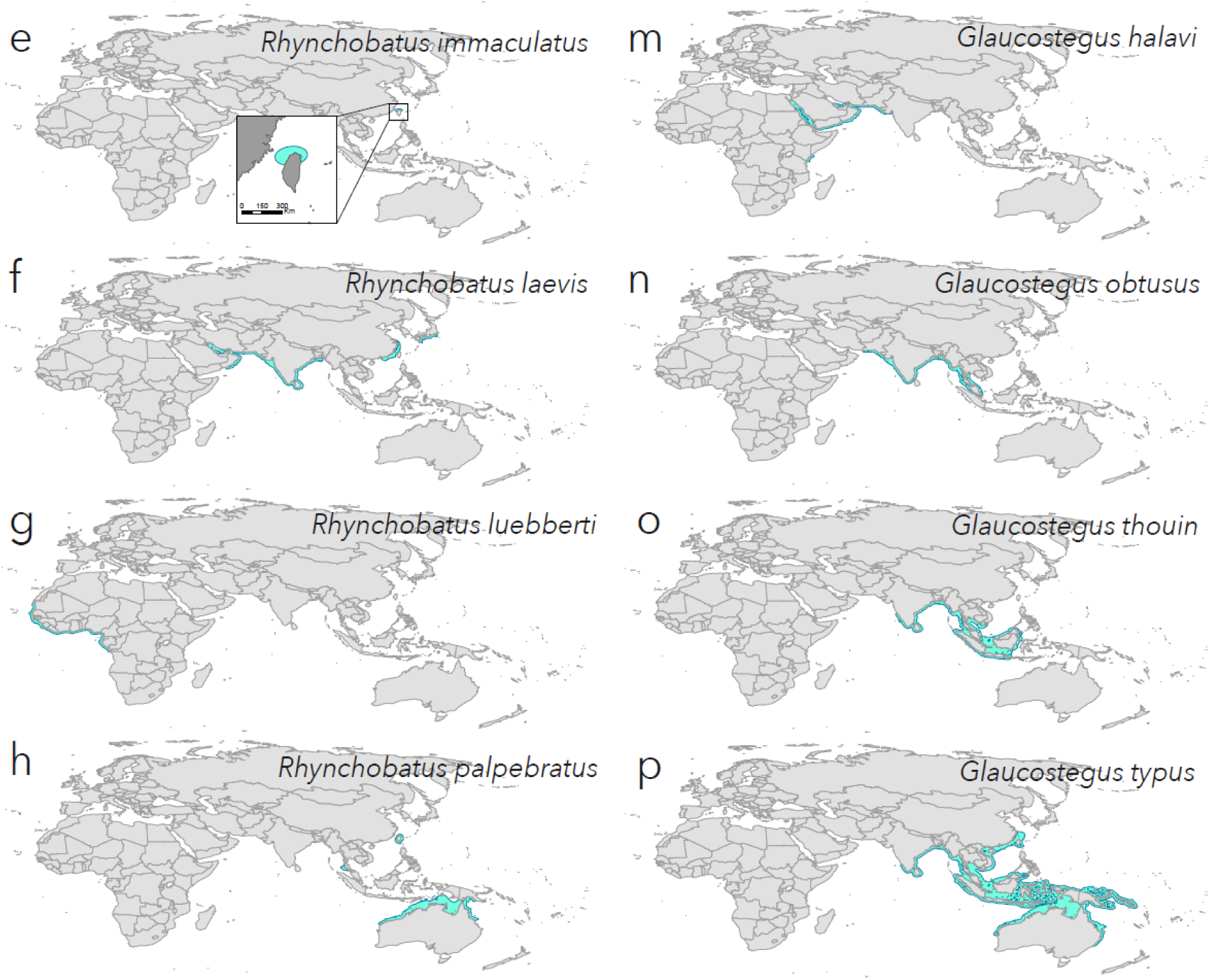
Individual species range maps for wedgefishes (a–j) and giant guitarfishes (k–p): (a) Rhina ancylostoma; (b) Rhynchobatus australiae; (c) Rhynchobatus cooki; (d) Rhynchobatus djiddensis; (e) Rhynchobatus immaculatus; (f) Rhynchobatus laevis; (g) Rhynchobatus luebberti; (h) Rhynchobatus palpebratus; (i) Rhynchobatus springeri; (j) Rhynchorhina mauritaniensis; (k) Glaucostegus cemiculus; (l) Glaucostegus granulatus; (m) Glaucostegus halavi; (n) Glaucostegus obtusus; (o) Glaucostegus thouin; (p) Glaucostegus typus.

## Notes

#### Summary of Updates

Revised Red List Indices; revised Rhynchobatus cooki status; revised Guinea management

